# A Systems-Level Framework for Drug Discovery Identifies Csf1R As A Novel Anti-Epileptic Drug Target

**DOI:** 10.1101/140087

**Authors:** Prashant K Srivastava, Jonathan van Eyll, Patrice Godard, Manuela Mazzuferi, Benedicte Danis, Catherine Vandenplas, Patrik Foerch, Karine Leclercq, Georges Mairet-Coello, Frederic Vanclef, Kirill Shkura, Liisi Laaniste, Andree Delahaye-Duriez, Rafal M Kaminski, Enrico Petretto, Michael R Johnson

**Affiliations:** Division of Brain Sciences, Imperial College Faculty of Medicine, London, UK.; New Medicines R&D, UCB Pharma, Braine-l’Alleud, Belgium; Clarivate Analytics (formerly the IP & Science business of Thomson Reuters), 5901 Priestly Dr., #200, Carlsbad, CA 92008, USA; Université Paris 13, Sorbonne Paris Cité, UFR de Santé, Médecine et Biologie Humaine; PROTECT, INSERM, Université Paris Diderot, Sorbonne Paris Cité; Duke-NUS Medical School, Singapore; MRC Clinical Sciences Centre, Imperial College Faculty of Medicine, London, UK

**Keywords:** Epilepsy, systems genetics, gene regulatory network, causal reasoning, drug discovery

## Abstract

The identification of mechanistically novel drug targets is highly challenging, particularly for diseases of the central nervous system. To address this problem we developed and experimentally validated a new computational approach to drug target identification that combines gene-regulatory information with a causal reasoning framework (“causal reasoning analytical framework for target discovery” – CRAFT). Starting from gene expression data, CRAFT provides a predictive functional genomics framework for identifying membrane receptors with a direction-specified influence over network expression. As proof-of-concept we applied CRAFT to epilepsy, and predicted the tyrosine kinase receptor Csf1R as a novel therapeutic target for epilepsy. The predicted therapeutic effect of Csf1R blockade was validated in two pre-clinical models of epilepsy using a small molecule inhibitor of Csf1R. These results suggest Csf1R blockade as a novel therapeutic strategy in epilepsy, and highlight CRAFT as a systems-level framework for predicting mechanistically new drugs and targets. CRAFT is applicable to disease settings other than epilepsy.

## INTRODUCTION

Despite advances in our understanding of disease processes at the molecular and cellular levels, modern drug discovery has failed to deliver improved rates of approval for mechanistically novel drugs^1^. One reason for the high rate of attrition in drug development, particularly for diseases of the central nervous system (CNS), is inadequate target validation in early stage drug discovery^1,2^. Optimism that advances in gene discovery would facilitate the validation of mechanistically novel targets has yet to materialize, and there is a requirement for new approaches to drug discovery.

Network-based systems analyses provide powerful new approaches to elucidate molecular processes and pathways underlying disease^3,4,5,6,7^. The power of the gene network approach comes from the analysis of multiple genes in functionally enriched pathways, as opposed to traditional single gene approaches that examine only one component of a complex system at a time. Using genome-wide transcriptional profiling in tissues relevant to the disease under investigation, gene co-expression network analysis can identify modules (i.e., sets of coexpressed genes) as candidate regulators and drivers of disease. Network-based drug discovery aims to harness this knowledge to identify drugs capable of restoring the expression of disease networks (modules) toward health^8,9^. At this systems-level framework, therapeutic compounds are judged not by their binding affinity to a particular protein, but by their ability to induce a transcriptional response (i.e., a gene expression profile) that is anti-correlated to the coordinated transcriptional program that underpins the disease state. This systems approach to disease modification is loosely termed the “signature reversion paradigm”, and is orthogonal to traditional concepts of drug discovery.

Currently, two broad approaches can be taken to identify drugs using the signature reversion paradigm. Firstly, drugs can be screened based on the chance overlap between a drug’s gene expression profile (drug-GEP) and a disease’s gene expression signature^10^. Whilst successful proof-of-concept examples of this approach have emerged^11,12^, the discovery of new drugs for diseases of the CNS using drug-GEP is constrained by a paucity of data relating to drug-induced transcriptional changes in cell types relevant to the CNS^13^. A second approach to leveraging gene networks to drug discovery has therefore emerged, which aims to map the underlying drivers and regulators of co-expression modules as candidate drug targets^7^. Successful examples of drug target discovery based on mapping the upstream regulators of disease modules include the identification of genetic drivers of disease modules using network expression quantitative trait loci (network eQTL) mapping^14,15^, and approaches that make use of regulatory interactions between transcription factors (TFs) and target genes (“interactomes”)^16^. As currently formulated however, these approaches have substantial limitations, in that the former may identify only large genomic regions in which several candidate genes could be equally implicated as master regulators of network expression, whilst the latter connects networks to TFs which in themselves usually lack chemical tractability as drug targets. Moreover, both methods have a requirement for sample-matched DNA variant and expression data which is often unavailable.

Given the current constraints to realizing the promises of network-based drug discovery we aimed to develop a new framework for drug target discovery. Our developed method termed “causal reasoning analytical framework for target discovery” (CRAFT) combines gene regulatory information with a causal reasoning framework to predict cell surface receptors with a direction specified influence on disease module activity. We specifically chose to develop a method connecting module expression to membrane receptors because more than half of all approved drugs target receptors^17^, thus maximizing the opportunity for drug repositioning and rapid experimental medicine proofs-of-principle. Although in this study we applied CRAFT to epilepsy, our method is equally applicable to any disease for which an underlying disease module can be identified.

Epilepsy is a highly debilitating disease of the CNS, primarily characterized by recurrent unprovoked epileptic seizures, but often associated with additional brain features including cognitive and behavioral impairments, and a heightened risk of death^18^. The causes of epilepsy can be broadly divided into cases that arise through no cause other than a genetic predisposition (“genetic epilepsy”), and epilepsy which develops secondary to an acquired brain injury such as following status epilepticus or head injury (“acquired epilepsy”)^19^. Whilst considerable advances have been made in identifying susceptibility variants for genetic epilepsy^20^, almost no progress has been made in identifying more effective antiepileptic drugs (AEDs)^21,22^. Indeed, currently, all available AEDs target a restricted number of putative mechanisms modulating transmission of nerve impulses at the synapse, and none of these drugs are disease modifying or curative^23^. Consequently, a third of people with epilepsy continue to have uncontrolled seizures despite all licensed AEDs, making improved therapy for epilepsy a global unmet need. Here, we set ourselves the challenge of discovering novel drugs and targets for acquired epilepsy. Although our *in vivo* pre-clinical validations focused on a single epilepsy network and its CRAFT-predicted regulator, our analyses highlighted several other potential new therapeutic targets for epilepsy.

## RESULTS

### Identification of candidate gene networks for epilepsy

A summary and description of the study workflow is shown in Fig 1. As a first step, we aimed to identify gene regulatory networks associated with epilepsy. To this end, we used an established post status epilepticus (SE) mouse model of acquired temporal lobe epilepsy (TLE)^24^. In this model of epilepsy the mice develop spontaneous recurrent seizures approximately 4 weeks after pilocarpine induced SE. As well as manifesting spontaneous epileptic seizures, these mice also reflect several of the behavioral and cognitive disturbances associated with TLE in humans and their response to AED therapy has been shown to be predictive of drug efficacy in human epilepsy^25^.

**Figure 1.**
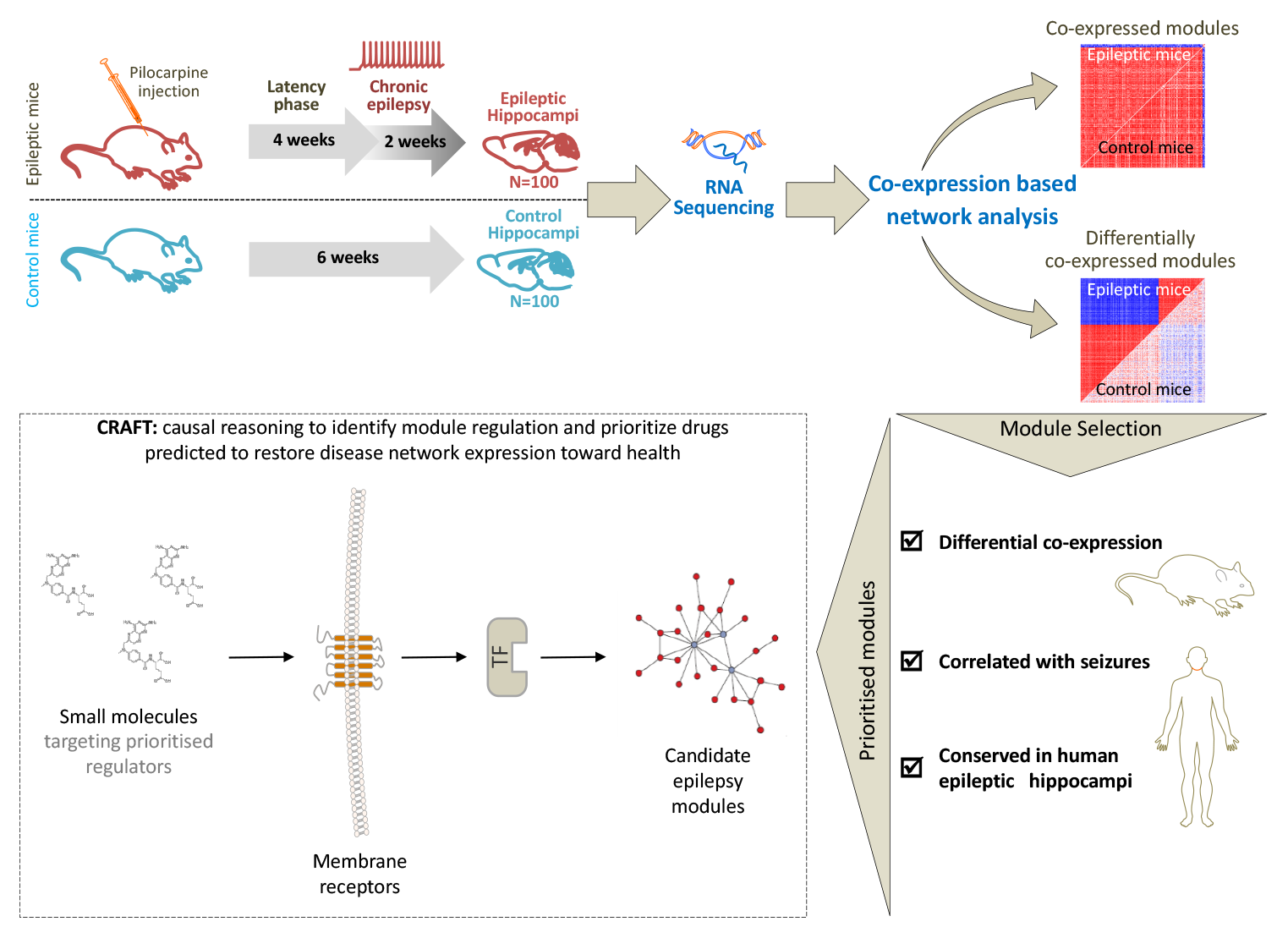
Experimental plan and study overview. We generated 100 mice with chronic epilepsy using the pilocarpine post-status epilepticus (SE) model of temporal lobe epilepsy (TLE). Controls were 100 healthy (i.e., pilocarpine naïve) matched littermate mice. At four weeks post SE each mouse was continuously video monitored for 14 days to record seizure frequency and severity. At the end of monitoring, each mouse was sacrificed and their hippocampus immediately frozen in liquid nitrogen. High throughput mRNA sequencing (RNA-seq) was generated using RNA from whole hippocampus samples from 100 epileptic and 100 control mice and the gene expression profiles were used to generate co-expression modules. Co-expression modules with a potential relationship to epilepsy were then prioritized using the following criteria: (*i*) differential co-expression between epileptic and healthy hippocampus in mouse and human TLE, (*ii*) correlation of module expression with seizure frequency (mouse), (*iii*) conservation in the human epileptic hippocampus. Coexpression modules meeting these criteria were considered candidate gene networks for epilepsy, and subjected to gene-regulatory analysis in a causal reasoning framework (CRAFT, see Text) to identify membrane receptors predicted to influence the expression of the disease-related network in a direction specified manner.

High-throughput sequencing of mRNA (RNA-seq) was performed on whole hippocampus samples from 100 epileptic mice and 100 control (i.e., pilocarpine naïve) mice (Methods). In total, 14,188 genes were expressed (log_2_FPKM>0) in at least 5% of samples and, of these, 9,013 genes showed significant (False Discovery Rate (FDR<0.05) differential expression (DE) between epileptic and healthy control mice (**Table S 1**).

To identify gene networks related to epilepsy, the set of genes expressed in the mouse epileptic hippocampus were first clustered according to their co-expression relationships (Methods). Briefly, Spearman’s rank correlation coefficients of expression were computed for all gene pairs and the pairwise correlation coefficients were used to perform hierarchical clustering based on Ward’s method^26^. The optimal number of co-expression modules was calculated using Elbow’s and pseudo F-index method^27^ (**Figure S.1**). This led to the identification of 29 modules consisting of 28 co-expression modules and an additional “module” (module 3) consisting of the un-clustered genes (see **Table S 2** for the full list of modules and their constituent genes). The 28 co-expression modules varied in size between 78 and 1,036 genes (mean and median module size was 255 and 188 genes respectively).

Analysis of the biological terms and canonical pathways enriched among the 28 coexpression modules in the mouse epileptic hippocampus revealed that the modules were generally enriched for specific functions - the top Gene Ontology (GO) biological processes enriched in each module are shown in **Figure S.2a** and the results of the functional enrichment analysis for each module are reported in full in **Table S 3**. Among the modules with overlapping functional terms, modules 5, 16 and 18 were enriched for “immune response” processes (Benjamini-Hochberg (BH)-corrected *P*=2.1×10^−11^, *P*=1.4×10^−6^ and *P*=1.3×10^−33^ respectively), and modules 10, 14, 26 and 29 were enriched for neuronal functions including “synaptic transmission” (BH *P*=4.4×10^−11^, *P*=0.02, *P*=4.0×10^−3^, and *P*=4.0×10^−4^ respectively).

To provide insights into the cell type expression of the modules we used cell type marker genes derived from single cell RNA-seq analysis of the mouse hippocampus (Methods)^28^. The individual modules demonstrated notable cell type specificity (**Figure S. 2b** and **Table S 4**). The cell type specificity of a module broadly corresponded to its functional enrichment – for example, “immune response” modules 16 and 18 were enriched for microglia marker genes whilst “synaptic transmission” modules 10, 14, 26 and 29 were specific for neuronal cell types.

To prioritize co-expression modules with a potential relationship to epilepsy we undertook a staged set of analyses summarized in **Figure S.3**. First, we tested if any of the modules were specific to the epileptic hippocampus using differential co-expression analysis. The differential co-expression paradigm postulates that a disease is linked to co-expression patterns (i.e., network topology, structure, interconnectivity) that are different in disease compared to healthy control states, potentially reflecting perturbed functional processes. Using methodology formulated by Choi and Kendziorski^29^ (Methods), 12 modules were found to be significantly (FDR<0.05) differentially co-expressed between the epileptic and control hippocampus, whilst 16 modules displayed conservation of co-expression (**Figure S.4** and **Table S 5a**). Of the 12 differentially co-expressed modules, modules 5, 16, and 18 were enriched for functional terms related to immune response processes, whilst modules 8, 10 and 21 were enriched for synaptic transmission and/or neuronal plasticity processes. As expected, the 16 modules with similar co-expression patterns in epileptic cases and controls were generally enriched for “housekeeping” functional terms unrelated to epilepsy such as cell morphogenesis and protein transport.

To further prioritize the modules in terms of their functional relationship to epilepsy, we explored the correlation between each module’s expression and seizure frequency. To this end we first quantified the frequency of behavioral seizures in each epileptic mouse by 14 days of continuous motion sensing 3D accelerometry synchronized with continuous video monitoring starting on day 28 post-SE (Methods). This revealed a diurnal variation of seizure occurrence in mice as well as clustering of seizures reflective of the classical patterns of human temporal lobe epilepsy (**Figure S. 5**)^30^. Next, using hippocampal gene expression data from each mouse, we summarized each module’s expression by its eigengene (i.e., its first principal component, PC1) and calculated the correlation between a module’s eigengene and seizure frequency (Methods). Of the 12 differentially co-expressed modules, nine modules (2, 5, 8, 10, 16, 18, 21, 22, and 24) had a pattern of expression that significantly (FDR<0.05) correlated with seizure frequency (Fig.2a).

**Figure 2.**
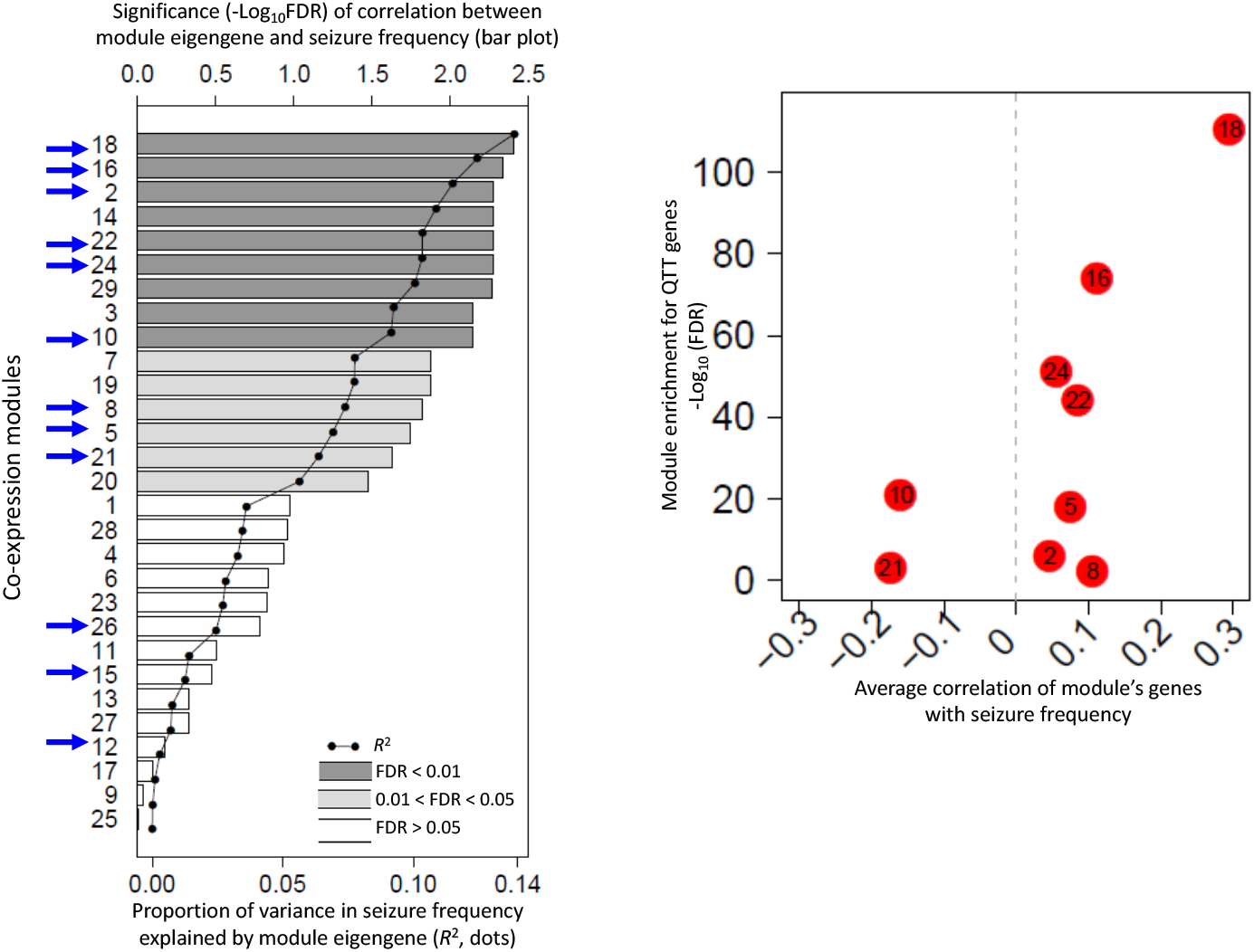
Correlation of module expression with epileptic seizures. **(a)** For each module identified in the epileptic mouse hippocampus, we plotted the significance (-Log_10_FDR) of the Spearman’s correlation between the module’s eigengene and seizure frequency (bar plot), and the percentage of variance in seizure frequency explained by the module’s eigengene (*R*^2^, dotted line). Modules marked with a blue arrow are the modules differentially co-expressed between the epileptic mouse hippocampus and the control mouse hippocampus. Modules highlighted in grey (bar plot) are significantly (FDR<0.05) correlated with seizure frequency. **(b)** Volcano plot of average (Spearman’s) correlation of a module genes with seizure frequency (X-axis) versus the significance of the module’s enrichment for genes individually correlated with seizure frequency (QTT genes) (Y-axis), for the 9 modules differentially coexpressed in epilepsy and correlated with seizures by module eigenegene.

To further explore the relationship between the expression of a module and seizures we correlated the expression of each individual gene in a module with seizure frequency using quantitative traits transcript (QTT) analysis^31^. Across all 29 modules, 1,833 genes had expression levels that significantly (FDR<0.05) correlated with seizure frequency (hereon termed “QTT genes”) (**Table S 6**). To assess the overall direction of correlation between a module and epileptic seizures, we plotted the average correlation of expression of genes in a module with seizure frequency against the module’s enrichment for QTT genes. Considering only the nine modules differentially co-expressed in epilepsy and correlated with seizures by module eigengene (i.e., modules 2, 5, 8, 10, 16, 18, 21, 22, and 24), module 18 (enriched for inflammatory processes and expressed in microglia) was the module most significantly positively correlated with seizures, whilst module 10 (synaptic transmission) was the module most significantly negatively correlated with seizures (Fig. 2b). The observed anti-correlation between down-regulated modules enriched in synaptic function and up-regulated modules enriched in inflammatory microglial pathways has recently also been described in autism spectrum disorder^32^.

For the nine modules differentially co-expressed in epilepsy and having a pattern of expressed that correlated with seizure frequency (i.e., modules 2, 5, 8, 10, 16, 18, 21, 22, and 24), we then assessed whether the module was conserved in the human epileptic hippocampus (Methods). Using human orthologues of mouse module genes and genome-wide gene expression data from 122 human epileptic hippocampus samples surgically ascertained from TLE patients^14^, we found that all nine modules were conserved in the human epileptic hippocampus (FDR<0.05) (**Table S 5b**). The conservation of these nine modules across human and mouse TLE provides an independent line of evidence for the validity of these modules, and further supports the relevance of the pilocarpine post SE mouse model of TLE to human TLE^14^.

As a final assessment of the relationship of these nine mouse TLE modules to human epilepsy, we tested whether each module was also differentially co-expressed in human TLE. In this analysis, for each module, we compared intra-module correlations in the human epileptic hippocampus with that in the non-diseased human hippocampus using post-mortem hippocampal samples ascertained from people with no history of psychiatric or neurological disease (Methods)^33^. Among the nine mouse modules differentially co-expressed in epilepsy and correlated with seizures, seven (5, 10, 16, 18, 21, 22, 24) were also differentially co-expressed in human TLE (**Table S 5b**). These seven modules were selected for further analysis. Specifically, we hypothesized that focusing on these seven modules (and by extension their enriched functional pathways) would provide a starting point for the development of new therapies for epilepsy.

Before proceeding to mapping the upstream regulators of these modules as candidate drug targets for epilepsy, since an important goal of our study was to identify mechanistically new drugs for epilepsy, we asked whether any of the seven modules could be considered to have a “known” relationship to epilepsy based on the published biomedical literature (Methods). Briefly, we first extracted published Abstracts for every gene in the genome using SCAIview webserver (www.scaiview.com) (3,811,179 abstracts with at least one gene-Abstract pair). The weight of evidence relating a particular gene to epilepsy was then quantified by determining if that gene’s co-citation with epilepsy (23,092 Abstracts with at least one gene-epilepsy co-citation) was more frequent than expected by chance (hypergeometric test, **Table S 7**). Then, by considering gene-epilepsy pairs significant at FDR<0.05, the modules were ranked according to their enrichment of gene-epilepsy pairs (hypergeometric test, **Table S 8**). Of the seven candidate epilepsy modules prioritized above, only module 10 (enriched for neuronal processes) was significantly (FDR<0.05) enriched for genes with a “known” relationship to epilepsy, suggesting these modules may be capturing novel functional relationships with epilepsy.

### Drug target prioritization through causal reasoning

The above analyses prioritize seven modules (5, 10, 16, 18, 21, 22, 24) as candidate modules for epilepsy by virtue of being (a) differentially co-expressed in both mouse and human TLE, (b) conserved in both mouse and human TLE, (c) have an expression profile that correlates with seizure frequency. From the pragmatic perspective of drug discovery, we set out to identify regulators of each of these modules as potential novel anti-epilepsy drug targets.

According to the signal reversion paradigm, if a module’s expression is related to the disease, then restoration of the module’s expression in the disease state toward health should be predictive of therapeutic benefit. We therefore set out to identify a drug-able target capable of restoring the activity of one or more candidate epilepsy module toward health. Since approximately 60% of existing drugs in clinical use target membrane receptors^17^, we decided to focus our search on finding membrane receptors exerting a regulatory influence over module activity. To this aim we developed and implemented a novel computational approach that combines ‘causal reasoning’ with gene regulatory information to ranks receptors based on the strength of their predicted effect on module expression (Methods). Briefly, using Clarivate Analytics MetaBase^®^ (version 6.15.62452), we first extracted information relating to known interactions between membrane receptors and transcription factors (TFs) via linear canonical pathways, and then between TFs and their target genes. To provide context to this “interactome”, only membrane receptors, TFs and target genes expressed in the mouse hippocampus were considered (resulting in a list of 1,624 expressed TFs and 307 expressed receptors). In a causal reasoning framework (logic summarized in Fig.3) there are multiple scenarios by which a membrane receptor can act via TFs on the set of genes in a module that are dysregulated in disease. For each of these scenarios the direction of effect of a membrane receptor on TFs and of the TFs on target genes is defined by a causal reasoning argument, which takes into account the directionality of the receptor>TF>target gene interactions and whether the network genes are over- or under-expressed in the disease state. For each scenario (Fig.3), the significance of the influence of a membrane receptor on a module’s gene expression can be predicted by considering the overlap between the direction-specified receptor effects on gene expression with the genes in a module that are over- or underexpressed in epilepsy (hypergeometric test; see Methods and **Figure S.6** for a full explanation of the method). This process allows membrane receptors to be ranked in terms of their predicted effect on module expression and the direction of that effect in terms of either activating or repressing the disease state, allowing the therapeutic directionality of receptor blockade or activation to be inferred.

**Figure 3.**
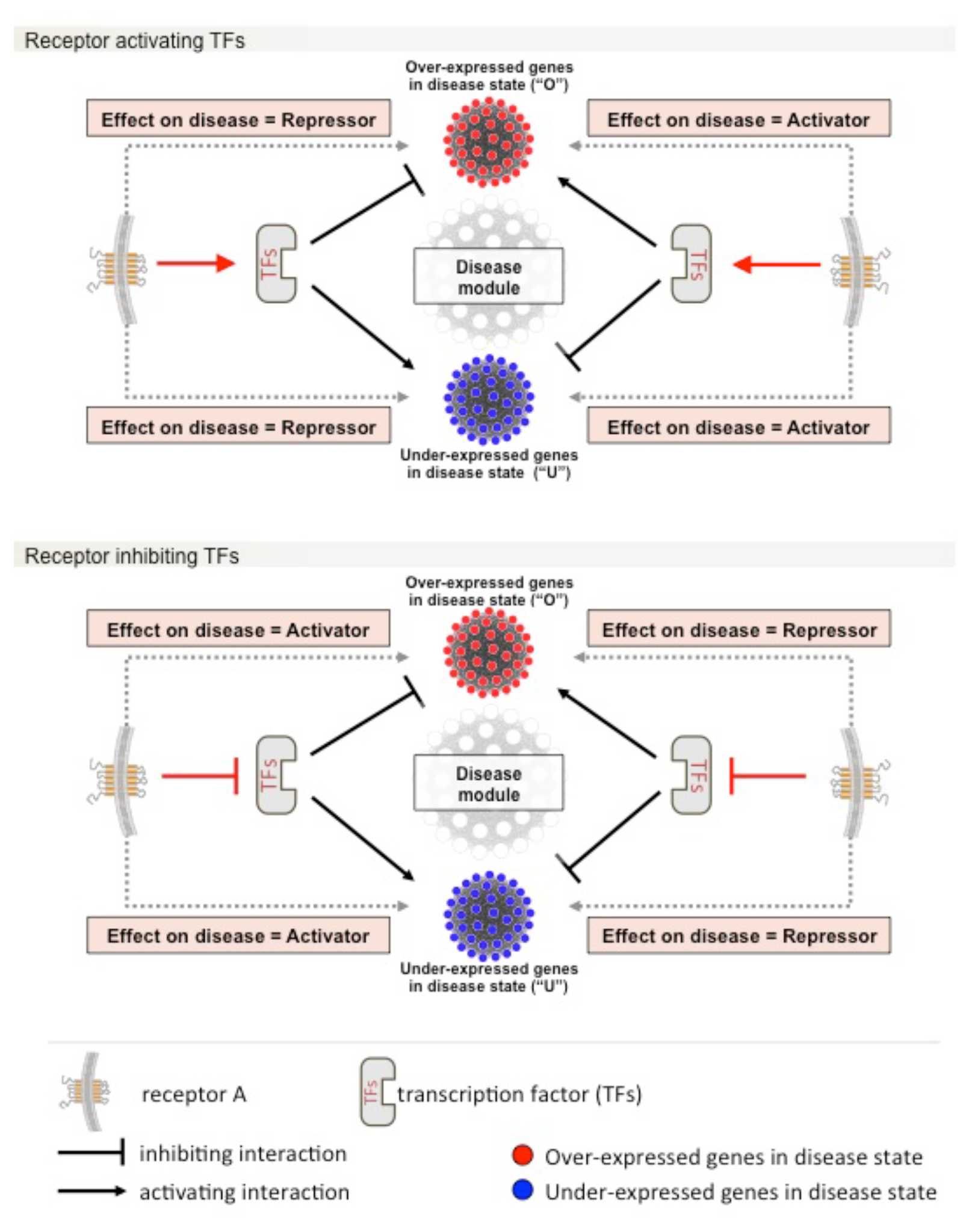
Causal reasoning framework. Logic for predicting magnitude, specificity and direction of effect of membrane receptors on the expression of genes in a disease module. A knowledge-based “interactome” connecting membrane receptors to module gene expression is first established based on experimentally validated connections between membrane receptors and transcription factors (TFs) in linear pathways, and between TFs and their target genes (genome-wide). This “interactome” is then integrated with information about whether the genes in the candidate module are over-(“O”) or under-(“U”) expressed in the disease state, allowing receptors to be classified as either disease “Activators” or disease “Repressors” which in turn permits the therapeutic directionality of receptor blockade or activation to be inferred. In the upper part of the figure we represent the positive activation of the TFs by the receptor “Receptor A”, whereas in the lower part of the figure we represent the opposite scenario of inactivation of the TFs by Receptor A. An illustrative example of the causal reasoning workflow is shown in **Figure S. 6**.

Of the seven candidate epilepsy modules, four (5, 16, 18, and 22) were significantly (FDR<0.05) enriched for one or more direction-specified receptor effect on module expression (**Table S.9a**). For each of these receptors, we plotted the proportion of genes in a module targeted by the receptor against the module’s–log_10_FDR enrichment for receptor (direction specified) target genes, allowing membrane receptors to be visualized in terms of their predicted directionality on the genes in a module which are over- or under-expressed in epilepsy, as well as the specificity and magnitude of the predicted effect (**Figures S.7a-d**). In support of the validity of our causal reasoning results, we found that membrane receptors related to interleukin-1 (IL-1) type 1 receptor/Toll-like receptor (IL-1R/TLR) signaling had a predicted direction of effect on epilepsy that was in agreement with the previously reported experimental evidence for that receptor^34^.

Of the many membrane receptors predicted to influence the expression of module genes in a direction specified manner, M-CSF receptor (also known as colony stimulating factor 1 receptor encoded by the *Csf1R* gene in the mouse) was predicted to be a regulator of two of the seven prioritized candidate epilepsy modules (modules 18 and 22). For both these modules, CRAFT predicted that Csf1R should “activate” the subset of genes in the module that are over-expressed in epilepsy. According to the CRAFT causal reasoning framework (Fig.3), Csf1R dependent activation of genes in modules 18 and 22 that are over-expressed in epilepsy should result in an activation of the disease state. By the same token, small molecule blockade of Csf1R should act to reduce the expression of these genes and the resulting restoration of the module’s expression toward health should be predictive of therapeutic benefit. Although there were several membrane receptors predicted to influence the expression of one or more candidate epilepsy module (**Table S.9a**), we chose to prioritize Csf1R for further analysis for the following reasons; (a) module 18 and 22 are enriched for functions related to inflammatory processes which have been previously linked to ictogenesis and epileptogenesis^35^, (b) Csf1R has not previously been linked to epilepsy (allowing for mechanistically new drug target discovery) and, (c) the availability of a tool compound, PLX3397, a known small molecule inhibitor of Csf1R^36^, by which we could experimentally test our causal reasoning predictions related to both module expression and impact on disease.

### Csf1R regulates module 18 genes

To test the predicted regulatory influence of Csf1R on modules 18 and 22 we first selected three genes in each module as markers of module expression (*Emr1, Aif1, Irf8, Gfap, ItgA5*, and *Serpine1*) on the basis that these genes (a) belonged to either module 18 or 22 and were among the set of genes predicted to be positively regulated by Csf1R, (b) are over-expressed in epileptic cases compared to controls, (c) are not predicted to be regulated by c-Kit (a kinase also inhibited by PLX3397)^37^. Healthy control mice were treated with PLX3397 at 3mg/kg/day or 30mg/kg/day or vehicle for 7 days (Methods). Hippocampus RNA was then extracted on day 7 and gene expression was measured by reverse transcriptase qPCR. Csf1R blockade with PLX3397 at 30mg/kg/day was associated with a significant decrease in the module 18 marker genes but not module 22 marker genes (Fig.4a). These results are consistent with Csf1R influencing the expression of module 18 (but not 22).

**Figure 4.**
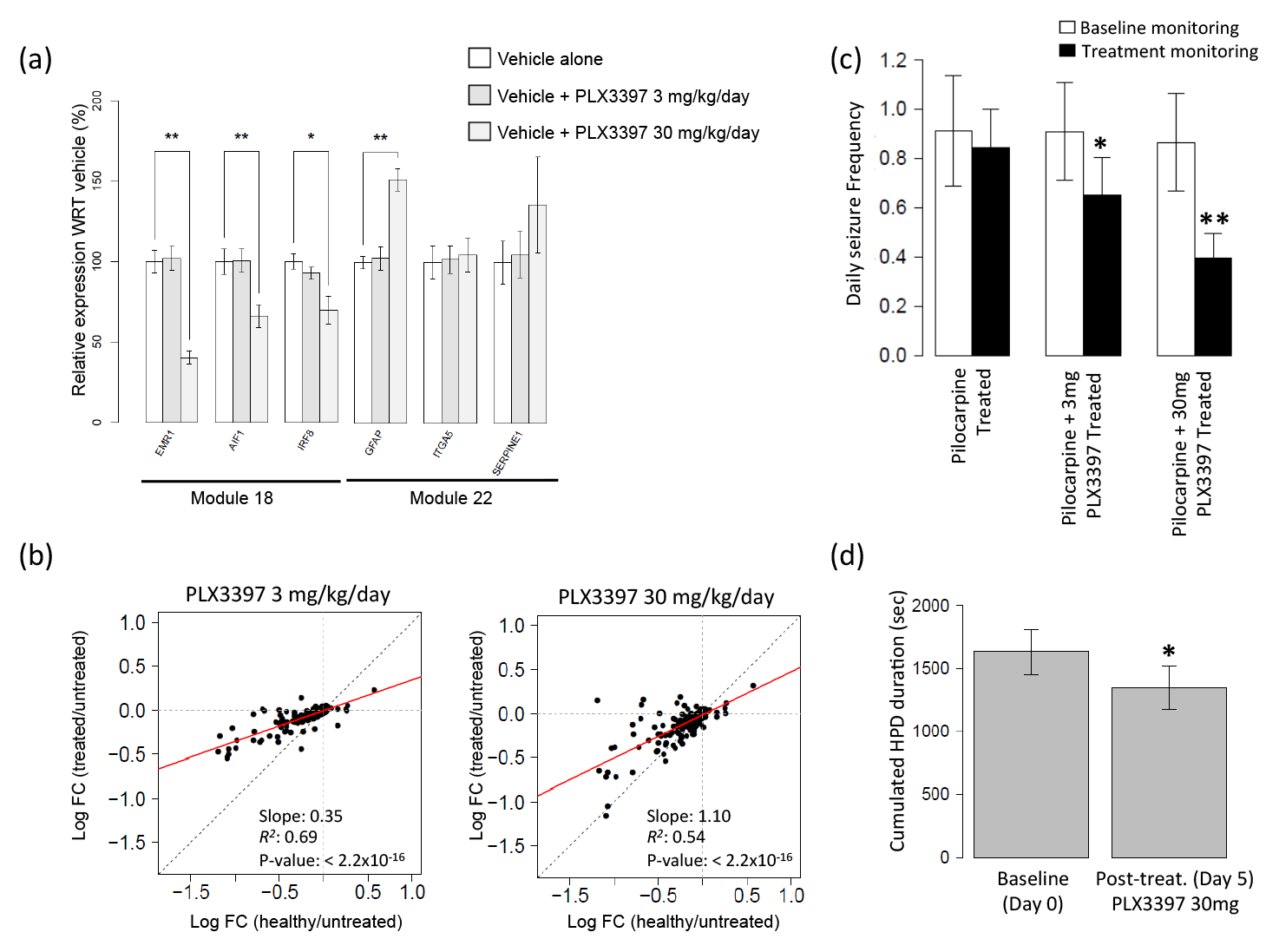
**(a) PLX3397 regulates Module 18 genes in control non-epileptic mice.** Pilocarpine naive mice were treated daily for 7 days with vehicle or PLX3397 at 3mg/kg/day or 30mg/kg/day. At the end of the treatment exposure, hippocampal RNA was extracted and the expression of marker genes in module 18 (predicted to be under the control of Csf1R) and marker genes in modules 12 and 22 (predicted not to be under the control of Csf1R) were assayed by reverse transcriptase qPCR. Consistent with the causal reasoning predictions, module 18 marker genes, but not marker genes from module 12 or 22, were significantly down-regulated by PLX3397 at 30mg/kg/day (*p-value<0.05, **p-value<0.01 – one tailed Welch’s t-test). **(b) Restoration of module 18 gene expression in epilepsy toward health by PLX3397**. Mice with epilepsy were treated with PLX3397 at 3mg/kg/day or 30mg/kg/day or vehicle alone. Hippocampal mRNA was extracted on day 14 of treatment and module 18 expression assayed by microarray. Red line indicates the linear negative correlation between the two conditions compared to a theoretical complete restoration of expression toward the healthy state (dotted black line). Treatment with PLX3397 resulted in a significant and dose-dependent (*P*=3.6x10^−14^) restoration of expression of module 18 genes toward health (toward the black diagonal). **(c) Efficacy of PLX3397 on seizures in the pilocarpine model of TLE**: Chronic epileptic mice were baseline-monitored for a week (white) before being daily administered with vehicle or PLX3397 at 3mg/kg/day or 30mg/kg/day and monitored for a second week (black). PLX3397 treatment induced a significant and dose-dependent decrease in daily seizure frequency (* p-value<0.05, ** p-value<0.01). **(d) Efficacy of PLX3397 on paroxysmal hippocampal discharges in the kainate model of epilepsy**. Kainate-induced chronic epileptic mice were EEG-monitored at baseline (day 0) for 2 hours prior to daily administration of PLX3397 at 30mg/kg/day for 4 days and then EEG-monitored on day 5 for 2 hours to assess drug efficacy. Treatment with PLX3397 led to a significant (* p-value<0.05) reduction in the duration of HPDs consistent with a therapeutic effect of PLX3397 in this model of pharmacoresistant epilepsy.

To confirm the regulatory influence of Csf1R on module 18, and to investigate the transcriptional response of module 18 to Csf1R blockade in more detail, we assayed the expression of all 171 genes in module 18 in response to PLX3397 treatment. Here, mice with epilepsy were treated with PLX3397 at 3mg/kg/day or 30mg/kg/day or vehicle alone. Hippocampal mRNA was extracted on day 14 of treatment and module 18 expression was assayed by microarray (Methods). In keeping with our causal reasoning predictions, we observed a significant (*P*<2.2x10^−16^) and dose dependent (*P*=3.6x10^−14^) restoration of module 18 expression toward health following treatment with PLX3397 (Fig.4b).

Although these results are consistent with a shift in expression of module 18 in epilepsy toward the healthy state by Csf1R blockade, PLX3397 has also been reported to deplete the brain microglial cell population as judged by Iba1 immunolabeling^36^, and module 18 is predicted to be highly expressed in microglia (**Figure S. 2b**). Using Iba1 immunolabeling we observed a similar decrease in Iba1 staining in the epileptic mouse cortex and hippocampus following treatment with PLX3397 (**Figure S.8**). However, Iba1 (encoded by *Aif1*) is both a component of module 18 and a predicted regulatory target of Csf1R and therefore Iba1 staining alone cannot distinguish between depletion of microglia by PLX3397 or a focussed down-regulation in *Aif1* gene expression following Csf1R blockade. To further explore these two possibilities, we tested all three microglia-expressed modules (i.e., modules 16, 18 and 24) for enrichment of genes down-regulated by PLX3397. Of the three modules, only module 18 was enriched (FDR<0.01) for genes down-regulated by PLX3397 (**Figure S.9**), suggesting a selective effect on module 18 expression by PLX3397 as opposed to an “apparent” down-regulation of module 18 genes secondary to gross microglial depletion. Consistent with this interpretation, we observed that the expression of the microglia marker *Sall1*^38^ was significantly up-regulated by PLX3397 (Log_2_ fold change = 0.17, FDR=2.16×10^−6^).

In addition to module 18’s enrichment for genes down-regulated by PLX3397 (**Figure S.9**), we also observed significant (P<0.01) enrichment of PLX3397 down-regulated genes in module 17 (functionally enriched for mitochondrial and metabolism related genes) and module 27 (functionally enriched for ribososme and RNA processing genes). Neither of these two modules are enriched for microglial marker genes (**Figure S.2b**), again consistent with an effect of PLX3397 on gene expression independent of gross microglial cell depletion. Re-inspection of our causal reasoning results (**Table S 9a**), revealed that both module 17 and 27 are predicted by the CRAFT framework to be regulated by Csf1R, albeit in a direction unspecified manner. As neither module 17 or 27 are differentially co-expressed in epilepsy nor correlated with seizures, PLX3397-induced change in the expression of modules 17 and 27 are not expected to impact the epileptic phenotype. However, we cannot exclude that PLX3397-induced changes in expression of these modules might contribute to potentially undesirable “off-target” adverse effects of PLX3397.

Taken together these analyses point to a restoration of the expression of module 18 in epilepsy toward health by PLX3397, mediated by Csf1R blockade. Under the signature reversion paradigm, if module 18 is a valid driver of epileptic seizures then PLX3397 treatment should exert a therapeutic effect in epilepsy. Whilst transcription factors are likely to be just one regulatory component of a module’s expression, and the extent to which a disease module’s expression needs to be shifted toward health in order for it to impact the clinical phenotype is unknown, we decided to test this prediction by investigating the therapeutic effect of PLX3397 in pre-clinical models of epilepsy.

### Csf1R blockade attenuates epileptic seizures in vivo

To assess the effect of PLX3397 on epileptic seizures, we first tested the effect of PLX3397 on epilepsy using the same pilocarpine model of TLE used for the discovery phase of this project (Methods). Mice in the chronic phase of epilepsy (i.e., exhibiting spontaneous recurrent seizures) underwent 14 days of continuous video monitoring as a baseline measurement of seizure frequency (“baseline period”). This was followed by daily treatment with PLX3397 (3mg/kg/day or 30 mg/kg/day) for 14 days with continuous video-monitoring of seizures during the treatment phase. As predicted by our causal reasoning framework, treatment with PLX3397 resulted in a significant (*P*<0.01) and dose-dependent decrease in the number of epileptic seizures (Fig.4c).

To confirm the therapeutic effect of PLX3397 on epileptic seizures, we repeated the analysis using a second independent pre-clinical model of epilepsy, the mouse intra-hippocampal kainate model of TLE (Methods)^39^. Here, seizure activity was assessed using continuous electroencephalographic (EEG) recordings^25^. The primary clinical outcome of this analysis was reduction in the duration of hippocampal paroxysmal discharges (HPDs) in response to PLX3397 treatment, which is considered a standard efficacy endpoint in this epilepsy model. In the kainate model, treatment with traditional AEDs usually only achieves a reduction in HPD duration at supra-therapeutic dosages and it has been suggested that this model of TLE is a model of drug-resistant epilepsy^40^. In this experiment, epileptic mice were EEG-monitored at baseline (day 0) for 2 hours prior to daily administration of PLX3397 at 30 mg/kg for 4 days and then EEG-monitored again on day 5 for 2 hours to assess drug efficacy. Treatment with PLX3397 led to a significant (*P*<0.05) reduction in the duration of HPDs consistent with a therapeutic effect of PLX3397 on epileptic seizures (Fig.4d) and replicating the results from the analysis of the pilocarpine model of TLE.

Notably, we observed that a single dose of PLX3397 (up to 30 mg/kg) did not display any anticonvulsant activity in acute models of epilepsy including either focal (6 Hz) or generalized seizure (maximal electroshock) models (data not shown). This distinguishes PLX3397 from all current AEDs which target neuronal excitability mechanisms, and supports a disease-context specific anticonvulsant effect of PLX3397 consistent with its measured effect on module 18 expression.

## DISCUSSION

In this study we used a network perspective of disease as a landscape for drug discovery. Under this framework, restoration of disease-related module expression toward health is predictive of therapeutic benefit, allowing “target” validation at the earliest stage of the drug discovery process. Based on these premises, we set out to develop and validate a novel predictive gene regulatory framework for drug discovery. Given the tractability of cell membrane receptors to drug development, and the large number of existing drugs targeting cell surface receptors, we specifically aimed to connect module expression to individual cell membrane receptors in a direction-specified manner, allowing the maximum opportunity for drug repositioning and rapid experimental proofs of principle.

Starting from genome-wide gene expression profiling of the epileptic mouse hippocampus, we first identified co-expression networks (modules) associated with the epileptic condition. The cell type specificity of these candidate modules and their functional processes were assessed using enrichment analyses, and the regulatory influence of cell membrane receptors over the selected modules was then inferred using gene-regulatory information in a causal reasoning framework (CRAFT).

Of the many cell surface receptors predicted to influence the expression one or more candidate epilepsy module in a direction-specified manner we chose to validate Csf1R for a number of reasons including (a) an absence of prior information connecting Csf1R to epilepsy (allowing for proof-of-concept that CRAFT can predict mechanistically novel targets) and (b) the availability of a tool compound (PLX3397) by which to test our causal reasoning predictions related to both Csf1R’s regulation of module 18 and, by extension, the relationship of module 18 to epilepsy. Analysis of module 18 expression using rtPCR quantification of marker gene expression was consistent with Csf1R influencing the expression of module 18, and subsequent analysis of transcriptional changes in the epileptic mouse brain in response to PLX3397 revealed a dose-dependent restoration of module 18 activity toward health by PLX3397. The predicted therapeutic effect of PLX3397 on epilepsy was then validated *in vivo* in two independent pre-clinical models of epilepsy, including a model of epilepsy resistant to standard AEDs. In addition to validating CRAFT as a predictive framework for drug target discovery, this result identifies Csf1R blockade as a novel therapeutic target in epilepsy and provides further evidence to support the role of innate immunity in the occurrence of seizures in acquired epilepsy^14,41^.

The ability to map the landscape of a disease in terms of its gene co-regulatory relationships (i.e., its interactome) and to combine this with direction-specified effects of membrane receptors on disease module activity offers considerable opportunities to accelerate the drug discovery process. Although we took advantage of experimentally validated interactions between TFs and target genes and between membrane receptors and TFs, meta-databases such as MetaBase^®^ have limitations in terms of the accuracy and completeness of this information, which places restrictions on the scope and accuracy of our current predictions. For example, the direction of effect of an interaction is often not specified in a database, and the relationship between membrane receptors and TFs is currently determined using linear pathways where knowledge is still incomplete. However, as the completeness of our knowledge of these regulatory relationships improves, including more detailed knowledge of cell-type specific interactions between TFs and target genes, so the accuracy and scope of CRAFT is expected to improve. At present, the major challenge was to establish proof-of-concept that disease networks combined with causal reasoning offer a valid framework to discovering mechanistically novel membrane receptors as drug targets, and this is what the framework described here makes possible. Although our causal reasoning framework was implemented using regulatory interactions from Clarivate Analytics MetaBase®, several other databases provide similar sources of information that can be adapted to the CRAFT framework. For example, biological pathway databases such as the Reactome pathway Knowledgebase^42^ and Pathway Commons^43^ provide well characterized linear signalling pathways which can be used to connect membrane receptors to transcription factors, whilst transcription factor target databases such as TRRUST^44^ provide information relating TFs to target genes. As well as having utility in prioritising novel drug targets for disease, CRAFT’s causal reasoning framework may ultimately have broader biological value in terms of understanding and modulating maladaptive transcriptional responses to environmental perturbations.

From a clinical perspective, our study identifies Csf1R as a novel drug target for acquired epilepsy, and highlights and further supports the potential for immunomodulatory therapies as a valid therapeutic approach in epilepsy^45^. Csf1R is a membrane receptor expressed by myeloid lineage cells including monocytes, macrophages and microglia^46^. It has been suggested that microglia are dependent on Csf1R signalling for their survival such that brain microglia are reported to be depleted from naïve mice following prolonged high dose treatment with PLX3397^36^. Although it could be argued that depletion of brain microglia might act to cause an “apparent” reduction in expression of genes in module 18 which are over-expressed in epilepsy, two other microglial modules were not significantly enriched for genes down-regulated by PLX3397 suggesting a more focused effect on module 18 expression according to our causal reasoning predictions. Additionally, we observed discordant changes in expression of microglial marker genes such as *Aif1* (which is a component of module 18 and predicted to be down-regulated by Csf1R blockade) and *Sall1*, which is not predicted to be a target of Csf1R and was not down-regulated by Csf1R blockade.

In conclusion, our approach (CRAFT) provides a gene regulatory and causal reasoning framework to identify membrane receptors as novel drug targets. As well revealing as Csf1R as a mechanistically novel target for acquired epilepsy, CRAFT highlighted many other regulators of candidate epileptic networks that may warrant further investigation as potential novel antiepilepsy drug targets.

## AUTHOR CONTRIBUTIONS

MRJ, RMK and EP conceived and designed the study and acquired funds. MRJ, RMK, EP PKS, JVE and PG, performed data analysis, and developed and implemented the methodology. Data generation and curation was performed by MM (lab), PKS, JVE, PG with contributions from KS, LL and ADD. MM, BD, CV, PF, KL, GM and FV carried out the laboratory experiments. RMK, EP, MRJ PKS and JVE wrote the manuscript. All authors read and approved of the final manuscript.

## ACKNOWLEDGEMENTS

We acknowledge funding from UCB Pharma, Imperial College NIHR Biomedical Research Centre (BRC) Scheme, the Singapore Ministry of Health and the European Union’s Seventh Framework Programme (FP7/2007-2013) under grant agreement number 602102 (EPITARGET, to E.P., and M.R.J.).

